# Large-scale analysis of SARS-CoV-2 spike-glycoprotein mutants demonstrates the need for continuous screening of virus isolates

**DOI:** 10.1101/2021.02.04.429765

**Authors:** Barbara Schrörs, Ranganath Gudimella, Thomas Bukur, Thomas Rösler, Martin Löwer, Ugur Sahin

## Abstract

Due to the widespread of the COVID-19 pandemic, the SARS-CoV-2 genome is evolving in diverse human populations. Several studies already reported different strains and an increase in the mutation rate. Particularly, mutations in SARS-CoV-2 spike-glycoprotein are of great interest as it mediates infection in human and recently approved mRNA vaccines are designed to induce immune responses against it.

We analyzed 146,917 SARS-CoV-2 genome assemblies and 2,393 NGS datasets from GISAID, NCBI Virus and NCBI SRA archives focusing on non-synonymous mutations in the spike protein.

Only around 13.8% of the samples contained the wild-type spike protein with no variation from the reference. Among the spike protein mutants, we confirmed a low mutation rate exhibiting less than 10 non-synonymous mutations in 99.98% of the analyzed sequences, but the mean and median number of spike protein mutations per sample increased over time. 2,592 distinct variants were found in total. The majority of the observed variants were recurrent, but only nine and 23 recurrent variants were found in at least 0.5% of the mutant genome assemblies and NGS samples, respectively. Further, we found high-confidence subclonal variants in about 15.1% of the NGS data sets with mutant spike protein, which might indicate co-infection with various SARS-CoV-2 strains and/or intra-host evolution. Lastly, some variants might have an effect on antibody binding or T-cell recognition.

These findings demonstrate the increasing importance of monitoring SARS-CoV-2 sequences for an early detection of variants that require adaptations in preventive and therapeutic strategies.

## Introduction

Since the first report of the severe acute respiratory syndrome coronavirus-2 (SARS-CoV-2) outbreak (1, 2), it has transformed into a global pandemic infecting and threatening death for millions of people all over the globe. By January 20, 2021, the World Health Organization (WHO) reported 94,124,612 confirmed cases and 2,034,527 deaths caused by the SARS-CoV-2 outbreak (3). On verge of the approval of SARS-CoV-2 vaccines which are designed to invoke immune responses against the spike-glycoprotein (spike protein), it becomes necessary to track the mutations in spike protein and study their relevance for current and upcoming vaccines. Also the recently approved neutralizing antibody bamlanivimab targets the spike protein of SARS-CoV-2 (4).

Subunits of the spike protein are valuable targets for vaccine design as the protein is responsible for viral binding and entry to host cells (5, 6). The spike protein consists of the N-terminal S1 and the C-terminal S2 subunits; the receptor-binding domain (RBD) in the S1 subunit binds to a receptor on the host cell surface and the S2 subunit fuses viral and host membranes (7). The receptor binding domain (RBD) of the SARS-CoV-2 spike protein recognizes human angiotensin-converting enzyme 2 (ACE2) as its entry receptor, similar to SARS-CoV (8). Interacting residues of the SARS-CoV-2 RBD with human ACE2 are highly conserved or share similar side chain properties with the SARS-CoV RBD (9). In addition, the SARS-CoV-2 RBD shows significantly higher binding affinity to ACE2 receptor compared to the SARS-CoV RBD. In order to repress the infection, blocking the RBD binding was effective in ACE2-expressing cells (5). Among the interacting sites in the SARS-CoV-2 RBD, particularly the amino acid residues L455, F486, Q493, S494, N501, and Y505 provide critical interactions with human ACE2 (10). These interacting residues vary due to natural selection in SARS-CoV-2 and other related coronaviruses (11). Similarly, worldwide SARS-CoV-2 genomic data shows ten RBD mutations which were caused due to natural selection by circulating among the human population (12). RBD mutations particularly at N501 may enhance the binding affinity between SARS-CoV-2 and human ACE2 significantly, improving viral infectivity and pathogenicity (10).

It is reported that continuous evolution of SARS-CoV-2 among the global population results into six major subtypes which involve the recurrent D614G mutation of the spike protein (13). Further, spread of such recurrent mutations within sub-populations might affect the severity of disease emergence and change the trajectory of the pandemic. Studies also report high intra-host diversity caused by low frequency subclonal mutations within a specific cohort (14). It is evident that changes in the SARS-CoV-2 genome over time might show new mutations which might influence the development efforts of of interventional strategies. The variability of epitopes of the RBD might hamper the development and use of neutralizing antibodies for cross-protective activities against mutant strains (15). Mutational variants of the spike protein might as well lead to escape variants with respect to pre-existing cross-reactive CD4+ T cell responses (16) or long-term protection from re-infection through T cell memory. Hence, there is a necessity of constant monitoring of the rapidly changing mutation rates in the spike protein in SARS-CoV-2, which could have significant impact on virus infection, transmissibility and pathogenicity in the current pandemic.

In this study, we gathered 147,413 genomic assemblies and 2,393 NGS sequencing datasets to detect non-synonymous spike protein mutations and infer their frequency within a given sample and the effect on potential antibody binding sites and known T cell epitopes.

## Methods

### SARS-CoV-2 assemblies

SARS-CoV-2 assemblies from human hosts were downloaded on October 2^nd^, 2020 from US National Center for Biotechnology Information (NCBI) Virus (protein sequences; 17) and on October 2^nd^, 2020 from GISAID (nucleotide sequences; 18). Pairwise alignments to the reference surface glycoprotein (NC_045512.2_cds_YP_009724390.1_3) were performed to extract the S gene sequences from GISAID samples using the R package Biostrings (version 2.52.0). Extracted sequences were translated with option if.fuzzy.codon = “solve”. Amino acid sequences of less than 100 length (440 samples) or premature stop codons (53 samples) were excluded from further analyses. Non-synonymous variants were determined by pairwise alignment (Biostrings, version 2.52.0) of the protein sequences to the translated reference sequence.

For three sequences obtained from NCBI Virus (accession IDs: QOE35701, QIQ50182, and QIQ50192), corresponding NGS data was available at the NCBI Sequence Read Archive (SRA, see section “NGS data processing”). Variant calling in the spike protein was in concordance between the assembly and the NGS data. Therefore, only the NGS data was used for further analysis.

### NGS data processing

All available NGS data for SARS-CoV-2 was downloaded on October 14^th^, 2020 from the NCBI SRA (https://www.ncbi.nlm.nih.gov/genbank/sars-cov-2-seqs/; 19) and filtered for whole genome fastq data from Illumina instruments with a human sample background. Data were aligned to the reference MN908947.3 (20).

Short-read whole genome sequencing data were aligned with bwa (version 0.7.17) mem (21). Output files in SAM format were sorted and converted to their binary form (BAM) using SAMtools (version 0.1.16) (22). Variants were retrieved from the alignment files using BCFtools (version 1.9) mpileup (http://samtools.github.io/bcftools/) with the options to recalculate per-base alignment quality on the fly, disabling the maximum per-file depth, and retention of anomalous read pairs. Variants in gene gp02 (i.e. S gene) were annotated using SNPeff (version 4.3t) “ann” (23).

### Filtering subclonal variants

NGS variants were filtered with at least 30 reads coverage and a fraction of supporting reads of at least 0.1 and less than 0.95 to identify high-confidence sub-clonal mutations (24).

### Calculation of solvent-accessible residues and corresponding solvent-accessible surface areas

Solvent-accessible residues of the spike protein were calculated using the rolling ball algorithm of the Swiss PDB Viewer (version 4.1.0; 25) with a parameter setting of >=30% accessible surface.

Solvent-accessible surface area (SASA) was calculated with tools from PyRosetta (version PyRosetta-4 2019) with default settings on reference pdb-structure “6vxx” for the spike protein (from PDB-Protein-Databank). SASA was calculated for every residue (in triplicates by the trimeric structure of the spike protein). The mutated structures were generated by introducing single mutations into the reference structure by tools from PyRosetta, too. This included merely a repacking of side-chains locally around the mutation side (with radius 3 Å), leaving the backbone unaltered.

### Published SARS-CoV-2 T-cell epitopes

SARS-CoV-2 antigens reported by Snyder et al. (26) where downloaded from https://clients.adaptivebiotech.com/pub/covid-2020 on 17NOV2020 (MIRA release 002.1).

## Results

### SARS-CoV-2 spike protein mutational profile from genome assemblies and NGS data

First, we determined the number of non-synonymous mutations in the spike protein per sample (for geographic background of the collected samples, see S1 Fig). Of the 146,917 analyzed genome assemblies (for exclusion of samples, see Methods section) and 2,393 NGS data sets, only 13.8% (20,246 samples) contained the WT spike protein (Fig 1A). Samples of mutant viruses exhibited only few mutations in the spike protein with less than ten mutations for all but 35 sequences. However, the mean and median number of mutations increased over time from December 2019 (mean: 0.14, median: to September 2020 (mean: 2, median: 2; Fig 1B). Overall, we detected 2,592 distinct non-synonymous mutations in the spike protein (Supplementary Table S1).

**Fig 1.**
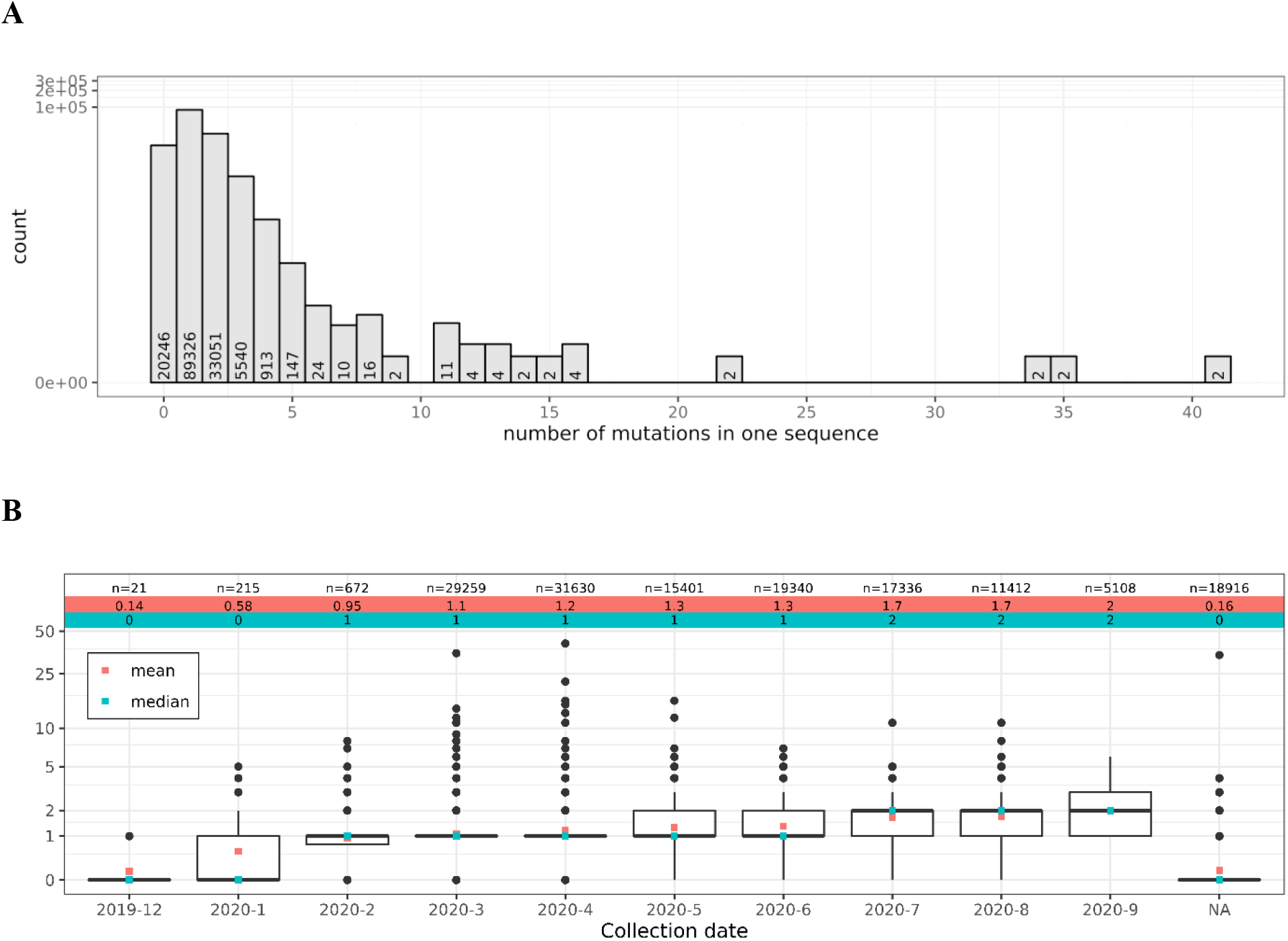
Most of the analyzed SARS-CoV-2 sequences differ from WT spike protein, but exhibit only few non-synonymous mutations. (A) The histogram shows the number of non-synonymous mutations in the spike protein detected in the analyzed samples. (B) The mean and median number of mutations per spike protein sequence increased over time.

### Recurrent variants in SARS-CoV-2 spike protein

Most of the observed variants in the assembly and NGS data sets were recurrent (Fig 2A) and only 33.2% of the variants were singular events in the combined assembly and the NGS data. The recurrent variants were distributed throughout the whole spike protein (Fig 2B, C). Among the recurrent variants, nine and 23 mutations were found in at least 0.5% of the mutant assembly and NGS samples, respectively (labeled variants in Fig 2 B, C). The most common mutation was D614G in both the genome assemblies (124,178 samples) and the NGS data (1,792 samples) located outside the RBD (positions 319-529), followed by the RBD variants S477N in the assemblies (11,483 samples) and N440K in the NGS data (75 samples). In total, 339 distinct mutations (227 recurrent) were detected in the RBD in the assemblies out of which only two were common to more than 0.5% of the mutated assembly sequences (Fig 2A). For the NGS samples, 61 mutations in total (24 recurrent) were found in the RBD (Fig 2B) and again only two were detected in at least 0.5% of the mutant NGS samples. Overall, 246 mutations were commonly found in the assembly and NGS data (Fig 2C).

**Fig 2.**
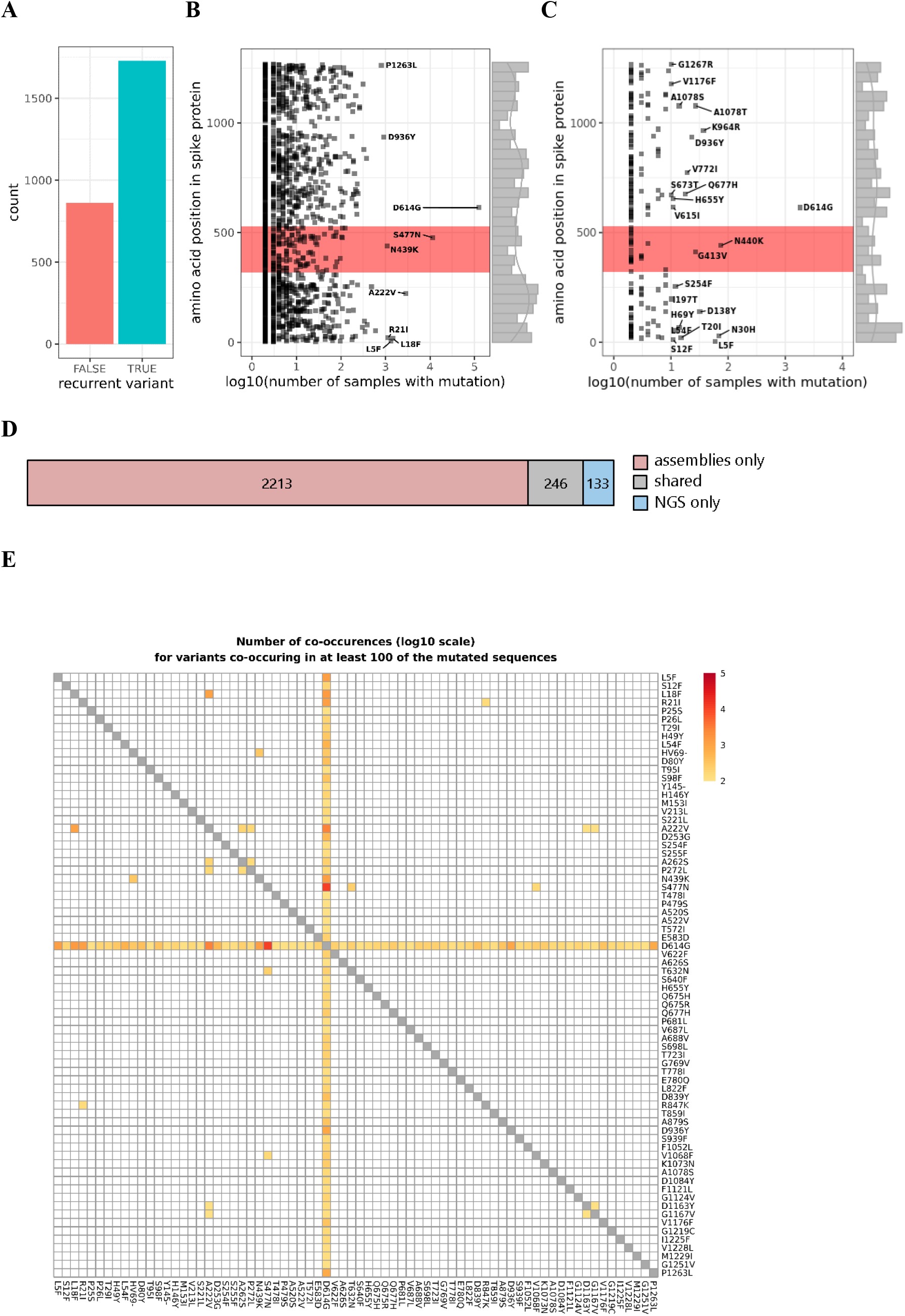
Recurrent variants are found throughout the whole spike protein. (A) Most of the detected variants were recurrent events occurring in at least two samples from the assembly or NGS data sets. (B, C) Each data point represents a distinct protein sequence mutation in the spike protein. The labels indicate the amino acid exchange for variants found in more than 0.5% of the assemblies (B) or NGS samples (C). The RBD is highlighted in red. (D) 246 variants (grey) were detected both in the assemblies and the NGS data. (E) A subset of 72 variants co-occurred in at least 100 of the mutated spike protein sequences (assemblies and NGS data combined). For better visibility, co-occurrences in less than 100 samples were set to 0 (white tiles).

Furthermore, 72 (2.8%) of the detected variants co-occurred frequently in at least 100 of the mutated spike protein sequences when we combined assembly and NGS data (Fig 2D). Most prominent here, was the variant D614G which was found in combination with 1,385 other variants. The combination S477N/D614G was detected in 11,470 samples. These represented the above mentioned two most frequent variants in the assembly data. The most frequent co-occurring mutations not involving D614G were L18F/A222V (1025 samples).

### Subclonal variants

In addition, we were interested in subclonal spike protein mutations (i.e. mutations with an observed variant frequency - as derived from the NGS reads - below 100%) which might either indicate co-infection with various SARS-CoV-2 strains and/or intra-host evolution of the virus. To this end, the fraction of variant supporting reads per sample of the detected mutations was determined. Most of the variants were observed with at least 95% of the reads supporting the respective variant nucleotide (Fig 3A, B). However, some mutations were only confirmed by a portion of the overlapping reads pointing to subclonal events. Filtering for a depth of at least 30 reads and a fraction of supporting reads between and 0.95 (24) resulted in 363 mutations observed in 292 samples (i.e. 15.1% of the NGS data sets with mutant spike protein) that could be classified as high-confident subclonal (Fig 3B). Most of these subclonal events were recurrent variants (Fig 3C). Especially in the earlier samples, but also in some later cases, the fractions of supporting reads within the same sample differed notably.

**Fig 3.**
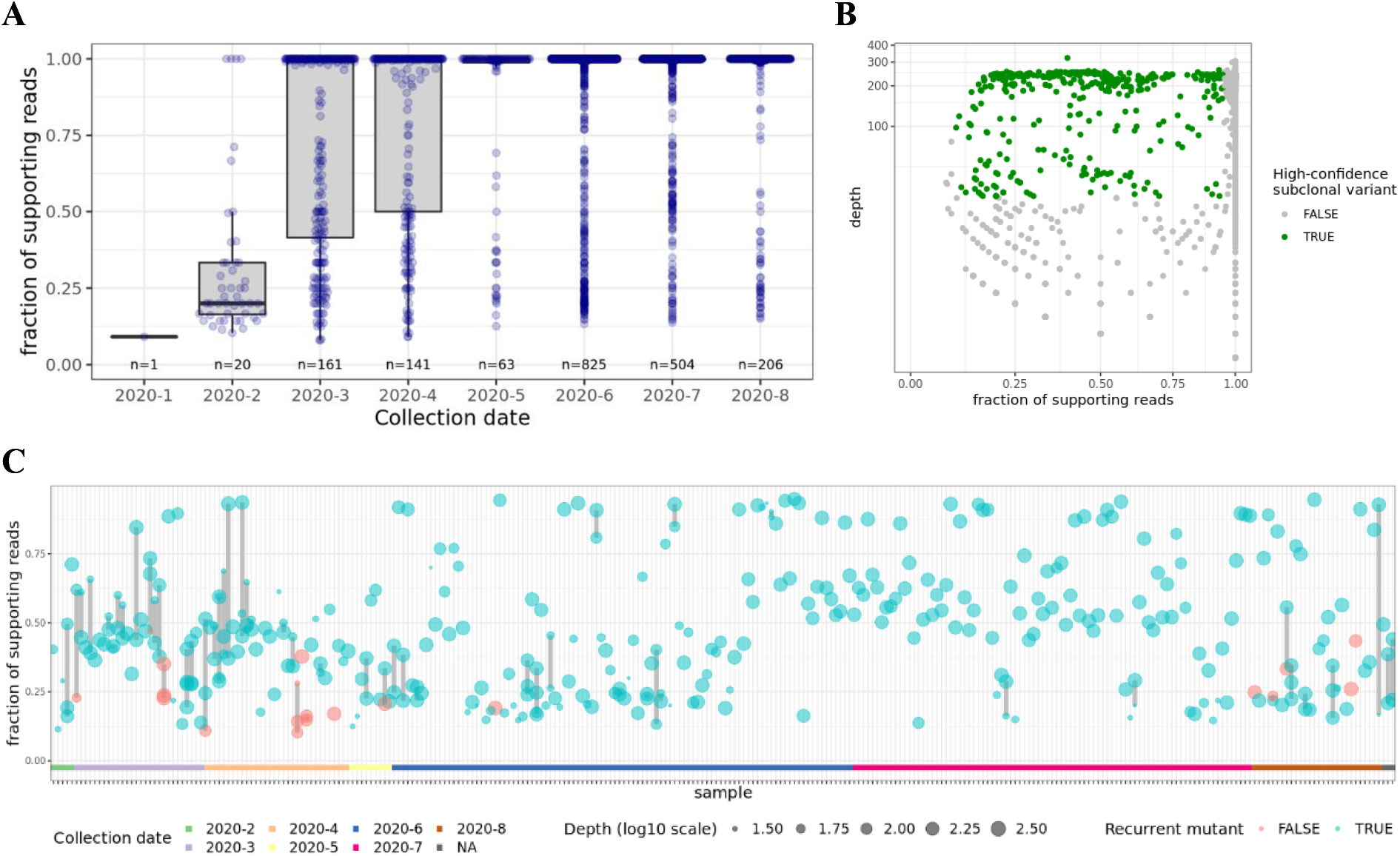
Variant frequencies of spike protein mutants indicate presence of multiple SARS-CoV-2 mutants in some samples. (A) The boxplot shows the distributions of the fraction of supporting reads of the mutations found in the NGS data. The numbers of underlying samples are indicated (n). Most of the observed variants have a variant allele frequency of >= 0.95 and can be accounted as clonal. (B) Filtering for high-confidence subclonal variants (green) with sequencing depth >= 30 reads and fractions of supporting reads between 0.1 and 0.95. (C) Sample-wise depiction of high-confidence subclonal events. Some of the observed subclonal variants were recurrent (blue) and only few were individual (red). The samples were ordered by collection date (see also color bar at the bottom of the plot) and point sizes indicate sequencing depth (log10 scale). Subclonal variants of the same sample are linked with grey lines. The fraction of supporting reads of variants found in the same sample differed notably in some cases.

### Effect of detected spike protein variants on potential antibody and T cell target sites

Next, we investigated whether the observed spike protein variants were relevant in the context of antibody binding or T cell recognition. In order to be visible for antibodies, a mutation has to hit a residue on the surface of the trimeric spike protein complex. 432 (16.7%) of 2,592 unique variants affected surface residues. For the 20 most frequent among these occurring in at least 50 samples, the change in SASA from wild type to mutation at the mutated residue position was investigated (Fig 4A). The SASA changed for all but one (H245Y) of the variants which might influence the accessibility of neutralizing antibodies. Furthermore, 2,544 (98.1%) of the 2,592 distinct variants hit at least one CD8+ or CD4+ T-cell epitope (Fig 4B) when compared to the T-cell epitopes reported by Snyder *et al.* (26) no matter if they were recurrent or individual events.

**Fig 4.**
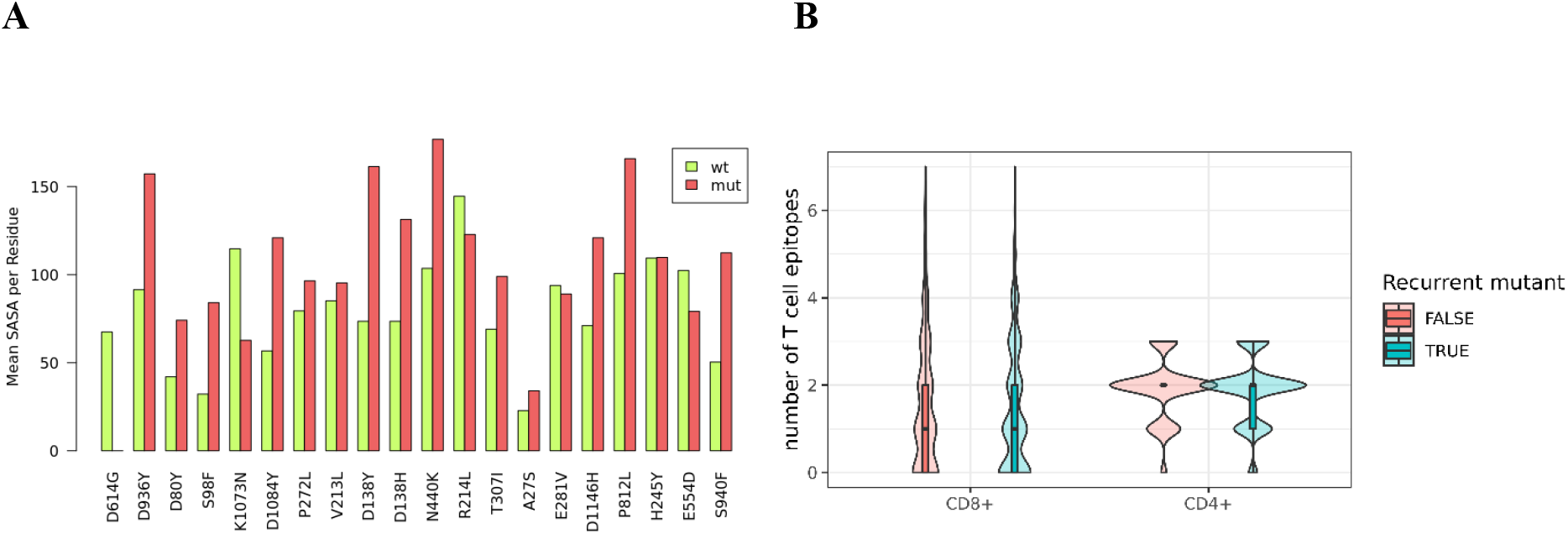
Variants affect antibody and T cell target sites. (A) Solvent-accessible surface area (SASA) values compared between wild type (wt) and mutation residue (mut) for surface variants occurring in at least 50 samples. The values are taken as the mean of the three replicated residues (3-meric structure of spike protein). Each time a new spike-protein structure has been generated by mutating the respective residue. The backbone of the mutated structure has not been re-modelled. The change in surface value is mainly due to change of amino acid and calculated optimal side-chain conformation. (B) The number of published T-cell epitopes (presented by MHC I or MHC II) that are affected by spike protein variants occurring in at least 50 analyzed samples is depicted. Most of the variants hit at least one epitope.

## Discussion

Our study sheds light on non-synonymous variants in the spike protein of SARS-CoV-2 in a large cohort of samples from all over the world. While most analyzed sequences vary from the reference sample from Wuhan, China, our analysis of almost 150,000 assembly and NGS samples shows an overall low mutation burden in the SARS-CoV-2 spike protein across different host populations (Fig 1). However, the mean and median number of variants per sample increased over time. Coronaviruses have fewer mutations compared to any other RNA virus due to its inherent 3’ to 5’ exoribonuclease activity (27). This suggests that the SARS-CoV-2 genome is genetically stable and the vast majority of mutations have no phenotypic effect such as virus transmissibility and virulence (28, 29). However, mutations of critical residues in the RBD of the spike protein might increase the virus transmission ability by enhancing the interaction (30). Furthermore, vaccines or treatments targeting the spike protein might become less efficient, if the number of variants in the spike protein increases further.

We identified a subset of mutations from the assembly and NGS data that are recurrent variants in the spike protein. Van Dorp *et al.* (31) have already reported such recurrent variants in SARS-CoV-2 evolution, which is a likely phenomenon of positive selection signifying the adaption of SARS-CoV-2 in human hosts. Furthermore, most recurrent variants show no evidence in increase of viral transmission and are likely induced by host immunity through RNA editing mechanisms (32). However, some variants might significantly influence SARS-CoV-2 transmission and infectivity. Among such variants, the non-synonymous D614G mutation has become most prevalent among several populations. We identified around 84.4% of the samples with a D614G variant, which supports a previous theory of an increasing frequency of the D614G variant in the global pandemic (30). Studies show evidence that the D614G variant is associated with high levels of viral RNA in COVID-19 patients, suggesting a role of D614G mutations in enhancing the viral infectivity in patients (30, 33–35). In contrast to these findings, it remains unclear whether the D614G variant makes the infections more severe or may impact vaccine design (36), as the viral load does not correlate with disease severity and the variant is not in the RBD of the spike protein, which interacts with the human ACE2 protein.

The RBD of the spike protein is a potential target for neutralizing antibodies and the variants in these regions might influence the infectivity and pathogenicity. We have identified high frequency variants in the RBD region from the assembly data, i.e. S477N, N439K, N440K and G413V (Fig 2B, C). S477N occurs frequently almost similar to the D614G variant and studies show that S477N has potential to affect the RBD stability and strengthen the binding with the human ACE2 protein (37, 38). In our study, S477N was most frequently co-occurring with D614G (Fig 2D). This combination was estimated to spread more rapidly than the D614G mutant alone (39). Other RBD variants such as N439K and N440K also show enhanced binding affinity to the human ACE2 receptor and result in immune escape from a panel of neutralizing monoclonal antibodies (40–42). Antibody-resistant RBD variants might affect the therapeutic potential of neutralizing monoclonal antibodies by escaping through disruption of epitopes. However, a significant portion of the detected variants represent individual events based on what could be deduced from the available data. This indicates the necessity to further collect SARS-CoV-2 isolates and monitor newly occurring variants. Here, the combination of assembly data (which appeared to be available in a timelier manner) and NGS samples (which also contain information on the clonality of the observed variants but which might be deposited with some delay) provide a valuable resource.

Further, we identified subclonal variants with a fraction of supporting reads between 0.1 and 0.95 at a sequencing depth of more than 30 reads in 15.1% of the NGS samples with mutant spike protein (Fig 3). Subclonal variants are indicative of within-host viral diversity leading to transmission of multiple strains (24). Low frequency variants could have been part of parallel evolution, where the same mutation rises to detectable frequencies in different lineages and it is observed as part of SARS-CoV-2 virus adaptation (43). Further, recurrent mutations might point to co-infection with multiple strains. Sample-specific variants in turn might rather indicate that the mutation occurred after infection within the host. This viral diversity within the host might prevent complete clearance after treatment and thus might lead to the development of resistant strains. Also, subclonal variants should be considered for vaccine design as these might represent the next generation of the virus.

The analyzed data sets also showed that a notable portion of the individual and recurrent mutations in the spike protein (98.1%) overlap with at least one known T-cell epitope. They also may change the solvent-accessible area and thus antibody binding when they involve surface residues of the trimeric spike protein complex as shown for the 20 most frequent solvent-accessible mutations. While we had no information on the HLA-restriction of the published T-cell epitopes, the influence on CD8+ T cell epitope generation by different HLA alleles was investigated for the three common mutations L5F, D614G and G1124V (44). These mutations were predicted to result in epitope gains, losses or higher or lower HLA binding affinities. Greaney et al. (45) presented a system to map mutations in the SARS-CoV-2 RBD that escape antibody binding. However, there is no overlap with our exemplary analysis on SASA changes. In agreement with the increase of the SASA of the mutation N440K, the binding affinity of this mutant to antibody REGN10933_REGN10987 is strengthened (46). All these findings demonstrate that SARS-CoV-2 mutants need to be set in the context of immune recognition to evaluate their implications for the global spreading of the pandemic and future preventive or therapeutic approaches in a timely manner.

## Conclusion and outlook

Human infections with SARS-CoV-2 are spreading globally since the beginning of 2020, necessitating preventive or therapeutic strategies and first steps towards an end to this pandemic were done with the approval of the first mRNA vaccines against SARS-CoV-2. Here, we show different types of variants (recurrent vs. individual, clonal vs. subclonal, hitting T-cell or antibody target sites vs. not-hitting) that can be incorporated in global efforts to sustainably prevent or treat infections. The underlying computational strategy might serve as a template for a platform to constantly analyze globally available sequencing data. In combination with a web-based platform to administer the results, this could help guiding global vaccine design efforts to overcome the threats of this pandemic.

The importance of our approach is underlined by the recently emerging UK lineage B.1.1.7 of SARS-CoV-2 (47), which is characterized by the accumulation of 17 variants; eight of those are located in the S protein. This lineage has a higher transmissibility compared to other lineages (48). The occurrence of this lineage questioned the efficacy of current vaccines, but first results showed that it at least unlikely will escape BNT162b-induced protection (49). Interestingly, the individual variants can be traced back to samples from March (P681H, T716I) and April (Y144del, N501Y, A570D) of 2020. It needs to be mentioned that the available data, although representing a large cohort, might not reflect the real distribution of the circulating variants as mostly samples of specific interest will be sequenced. International sequencing efforts, combined data analysis and prediction of variant impact will be important tools for the future in order to ensure an early detection of such genomic variants of concern.

## Acknowledgments

We thank Pablo Riesgo Ferreiro and Patrick Sorn for critical discussions. We gratefully acknowledge the authors from the originating laboratories responsible for obtaining the specimens, as well as the submitting laboratories where the sequence data were generated and shared via GISAID, NCBI Virus or the NCBI SRA, on which this research is based.

## Author contributions

Conceptualization, U.S., M.L., and B.S.; Formal Analysis, B.S., R.G., T.B., and T.R.; Investigation, B.S., R.G., T.B., and M.L.; Writing – Original Draft, B.S., R.G., and T.B.; Writing – Review & Editing, B.S., R.G., U.S., and M.L.

## Supporting information

**S1 Fig.**
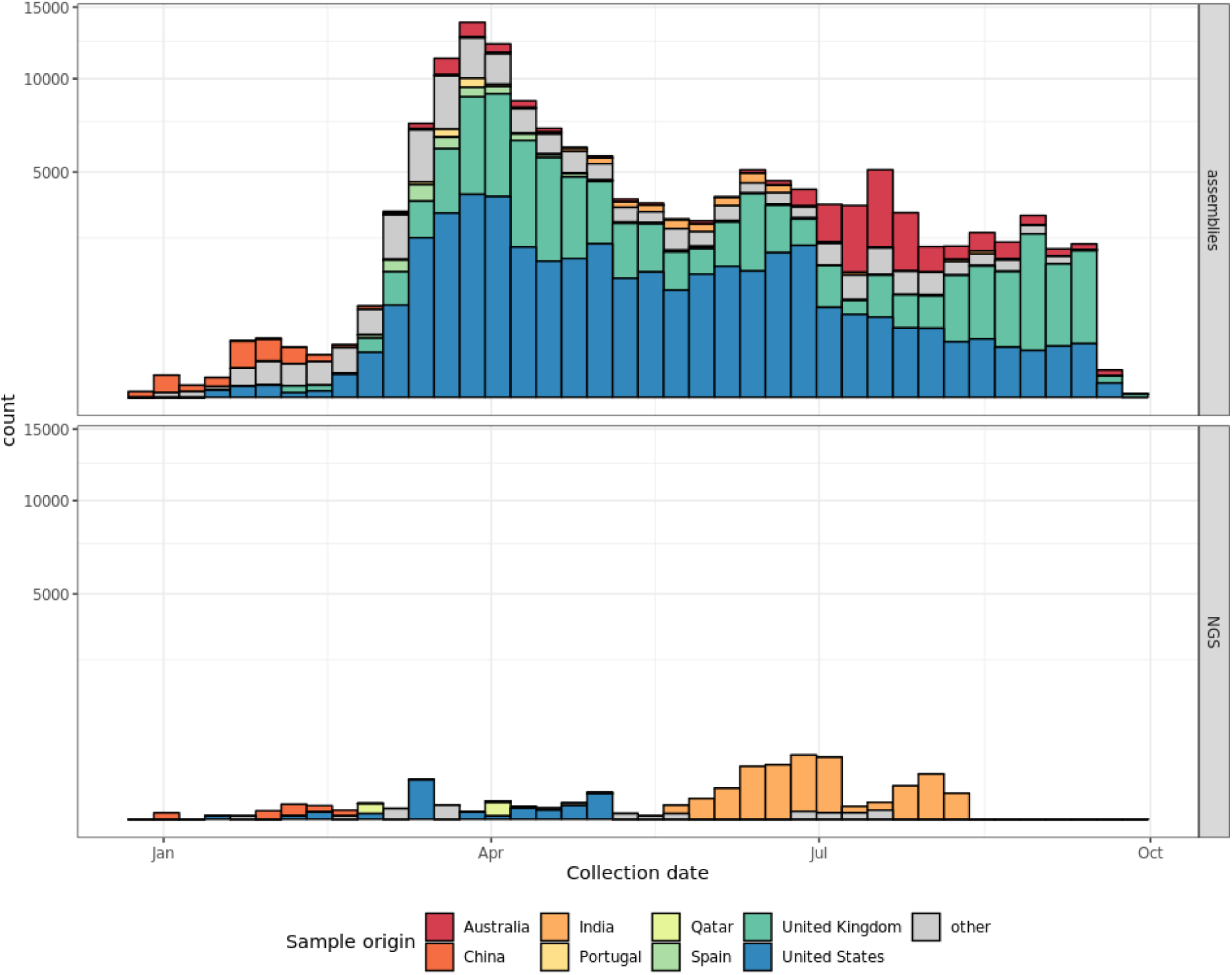
Number and origin of publicly available SARS-CoV-2 sequence data over time. The histogram shows the number of SARS-CoV-2 assembly sequences deposited at GISAID and NCBI Virus and NGS data deposited at SRA as of 02OCT2020. Color coding indicates the sample origin. Countries summarized as “other” include: Algeria, Andorra, Argentina, Aruba, Austria, Bahrain, Bangladesh, Belgium, Belize, Benin, Bosnia and Herzegovina, Botswana, Brazil, Brunei, Bulgaria, Cambodia, Canada, Chile, Colombia, Congo [DRC], Costa Rica, Crimea, Croatia, Cuba, Curacao, Cyprus, Czech Republic, Denmark, Dominican Republic, Ecuador, Egypt, Faroe Islands, Finland, France, Gabon, Gambia, Georgia, Germany, Ghana, Gibraltar, Greece, Guam, Guatemala, Hong Kong, Hungary, Iceland, Indonesia, Iran, Iraq, Ireland, Israel, Italy, Jamaica, Japan, Jordan, Kazakhstan, Kenya, Kuwait, Latvia, Lebanon, Lithuania, Luxembourg, Madagascar, Malaysia, Mali, Mexico, Moldova, Mongolia, Montenegro, Morocco, Myanmar, Nepal, Netherlands, New Zealand, Nigeria, North Macedonia, Norway, Oman, Pakistan, Panama, Peru, Philippines, Poland, Puerto Rico, Reunion, Romania, Romania, Russia, Saudi Arabia, Senegal, Serbia, Sierra Leone, Singapore, Slovakia, Slovenia, South Africa, South Korea, Sri Lanka, Suriname, Sweden, Switzerland, Taiwan, Thailand, Timor-Leste, Tunisia, Turkey, Uganda, Ukraine, United Arab Emirates, Uruguay, Venezuela, Vietnam, Zambia and unknown.

**S1 Table. Overview of the 2,592 distinct non-synonymous mutations in the spike protein of SARS-CoV-2 detected in genome assemblies and NGS data sets.**

